# Real-world testing of the durability claims of a commercially available spray-on surface biocide

**DOI:** 10.1101/2025.04.17.649382

**Authors:** Shilpa Saseendran Nair, Alana Cavadino, Simon Swift, Siouxsie Wiles

**Affiliations:** Department of Molecular Medicine and Pathology, Waipapa Taumata Rau University of Auckland, Auckland, New Zealand; Department of Epidemiology and Biostatistics, Waipapa Taumata Rau University of Auckland, Auckland, New Zealand

## Abstract

Self-disinfecting biocidal surfaces have been proposed as a solution to prevent infections caused by the transmission of microorganisms from frequently touched surfaces in hospitals and other settings. For example, surface anchoring quaternary ammonium salt (SAQAS)-based biocides are purported to maintain their antimicrobial potency for up to 30 days and are available as spray-on formulations. We tested the 30-day potency claim of a commercially available spray-on SAQAS-based biocide (“SAQAS-A”) under “real-world” conditions in a microbiology laboratory where regular cleaning is routine. To do this, we determined the background microbial burden on high-traffic floor areas, routinely used bench areas, frequently touched handles, and glass surfaces for 30 days before and after applying SAQAS-A. Experiments were repeated three times, and data was analysed using two Generalised Linear Mixed Models. We observed that for >80% of bench samples, the number of viable bacteria recovered was below the highest acceptable level of 2.5 CFU/cm^2^, with minimal reduction in CFU recovery observed after SAQAS-A treatment. In contrast, the number of floor and glass samples in which the microbial burden exceeded 2.5 CFU/cm^2^ was greater after biocide application (12.7 and 73%, respectively) than before (4.8 and 37%, respectively). Analysis of all data using two statistical models confirmed that the application of SAQAS-A had no antimicrobial effect. In conclusion, our results indicate that SAQAS-A was ineffective in preventing surface contamination over 30 days in a real-world scenario where routine cleaning occurs.

**Importance:** We tested the claim that a commercially available spray-on biocide could protect surfaces from microbial contamination for 30 days. In the presence of routine disinfection, we found that spray-on applications of the surface anchoring quaternary ammonium salt-based biocide did not protect frequently touched surfaces in a working microbiology research lab for up to 30 days. Our data show that users of this type of product should be aware of the limitations of manufacturers’ claims of “continued protection”, and manufacturers should consider formulations that more reliably attach to surfaces. Similarly, regulatory and consumer protection agencies should provide clear guidance for companies wishing to make durability and longevity claims for biocidal surface applications.

## Introduction

It is now well established that surfaces play a role in the transmission of pathogens to patients within hospitals(1, 2). Frequently touched surfaces such as medical equipment and items in the hospital environment (for example, bed rails) act as a reservoir for bacteria and viruses(3, 4) and can lead to hospital-acquired infections (HAI) with severe outcomes(5–7). So, while environmental transmission via fomites is a problem, it also presents an opportunity for control if reservoirs can be eradicated or routes of transmission broken. Environmental cleaning and surface disinfection are important hygiene measures that contribute to minimising the transmission of hospital-acquired pathogens(8, 9). However, there are several drawbacks associated with currently available strategies. These include variations in bactericidal efficacy, microbial resistance, cost, non-eco-friendly chemical constituents, and the likelihood of recontamination. Standard operating procedures for cleaning and decontamination also differ within and between hospitals(9). These drawbacks have driven the development of alternative technologies, such as self-disinfecting biocidal surfaces(10, 11).

Ideally, biocidal surfaces should exhibit durable, broad-spectrum antimicrobial activity and be made from biocides that are biodegradable and non-toxic to humans and the environment, thereby avoiding long-term environmental persistence. Surface design ideas can be classified into four groups(12, 13): (i) antifouling surfaces where modifications prevent bacterial attachment; (ii) anti-bacterial surfaces where biocidal agents are incorporated into materials and leach from the surface to kill bacteria that come close enough to attach; (iii) anti-bacterial surfaces where biocidal agents are anchored to the surface and kill bacteria on contact; and (iv) anti-bacterial surfaces with modifications that combine the advantages of anti-fouling, biocide releasing, and/or contact killing to develop dual-function surfaces.

Contact killing at surfaces through the application of antimicrobial polymers is receiving attention. The advantages of this type of surface biocide include the retention of the antimicrobial agent for durable killing activity without leaching into the environment and the requirement of a small amount of the active compound in a thin surface layer for eco-friendly activity(14). The polymers can either be incorporated into the material or applied onto the material surface(14–16). The covalent bonding of these polymers to the surface can help to retain the polymer at the surface, giving durable activity. However, limitations include the durability of coatings on different surfaces, the emergence of resistant strains after long periods of continuous exposure, and microorganisms associated with particles not coming into lethal intimate contact with the biocide(17, 18).

The antimicrobial activity of 3-(trimethoxysilyl)-propyldimethyloctadecyl ammonium chloride, an organosilicon quaternary ammonium compound, has been commercialised for use as a surface sanitiser. The commercial products are manufactured in spray formulations that are sold ready to use at room temperature(19). Marketing materials claim that these products exhibit broad-spectrum antimicrobial activity, with durability and stability, and are non-toxic to human cells, making them eco-friendly. According to the manufacturers, when applied, these biocides become covalently attached to the surface and polymerise to form a stable and long lasting coating(19). After application, the biocide dries onto the surface and is proposed to form a thin coating of molecular spike-like structures at the surface, which electrostatically attract microorganisms, killing them by rupturing the cell envelope. Hence, it is argued that the possibility of forming resistant strains is reduced because this compound kills mechanically. In some cases, it is claimed that the biocide is stable and can ensure protection for around one month, preventing recontamination in that period(20).

Introducing antimicrobial surfaces aims to reduce the microbial burden on surfaces and any subsequent transmission of pathogens. The success of this approach should be determined by employing the surface in a real-life environment(21, 22). Several studies have shown the effectiveness of copper-surface interventions in hospitals by replacing the surfaces with copper or copper alloys(23–26). Microbial burdens fell below the generally acceptable burden of environmental bacteria on hospital surfaces of 2.5 colony forming units (CFU)/cm^2^ (24, 27, 28). Copper films were also found to be viricidal when used for 90 days within high touch locations of an active metropolitan bus and railcar(29). Other studies have established the potential of N-halamine and photocatalytic coatings that release antimicrobial chlorine and reactive oxygen species, respectively, and can be recharged by exposure to either dilute bleach or the appropriate wavelength of light(30, 31). Several studies have also evaluated the efficacy of quaternary ammonium compound sprays, but typically only for a short period after application (for example, 2-24 h)(32–34).

We have previously shown that a commercially available SAQAS-type spray-on biocide protects treated glass and low-density polyethylene surfaces against model Gram-positive and Gram-negative species and antibiotic-resistant isolates(35). The manufacturers of spray-on SAQAS products have claimed that treated surfaces will retain activity for up to 30 days after application, and regular cleaning does not interfere with the activity of the treated surface. In this study, we test these claims in a Physical Containment (PC) 2 microbiology laboratory, equivalent to Biosafety Level 2 (BSL-2), as a surrogate for the hospital environment. The designated laboratory areas were regularly swabbed to determine the background microbial load over a 30-day period. A commercial SAQAS product (“SAQAS-A”) was then applied according to the on-label instructions, and the surfaces were swabbed for plate counts to examine any changes in the microbial burden over a further 30-day period. The generally acceptable burden of environmental bacteria on hospital surfaces is 2.5 CFU/cm^2^, and this value is used as a measure of the success of the surface treatment(24, 27, 28).

## Results

### SAQAS-A did not offer an advantage as a biocidal treatment for laboratory benchtops or door glass

The microbial burden present on laboratory benches and door glass was determined before and after SAQAS-A application (Fig.1). The data show that for the majority of samples, the number of viable bacteria recovered was below the highest acceptable level of 2.5 CFU/cm^2^, with minimal reduction in CFU recovery observed after biocide treatment. For laboratory benches, 12/63 (19%) of samples (5/21 [Trial 1], 3/21 [Trial 2], and 4/21 [Trial 3]) exceeded the acceptable level before treatment, while 7/63 (11%) of samples (2/21 [Trial 1], 1/21 [Trial 2], and 4/21 [Trial 3]) exceeded the acceptable level after treatment (Fig.1 A/C/E). Similarly, 3/63 (4.8%) of door glass samples (0/21 [Trial 1], 2/21 [Trial 2], and 1/21 [Trial 3]) exceeded the acceptable level before treatment, while 8/63 (12.7%) samples (1/21 [Trial 1], 3/21 [Trial 2], and 4/21 [Trial 3]) exceeded the acceptable level after treatment (Fig.1 B/D/F). The laboratory standard operating procedure requires benches to be disinfected after use. Our data suggest that the application of a biocidal SAQAS treatment does not offer any additional advantage for surfaces that are routinely disinfected, such as laboratory benchtops, or those which are infrequently contaminated.

**Figure 1.**
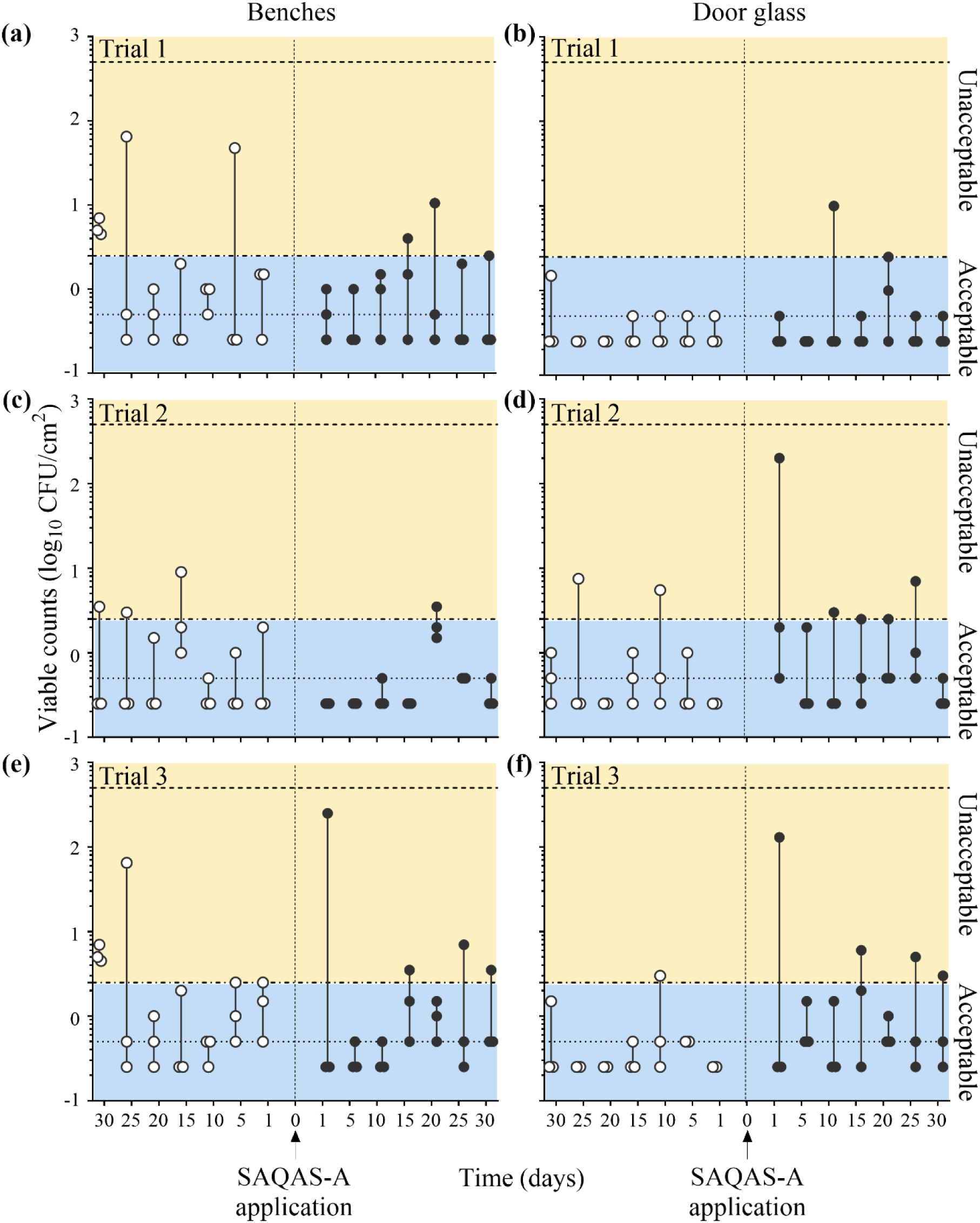
Bacterial counts recovered from benches and door glass before and after the application of SAQAS-A. Data presented as individual viable counts (log_10_ colony forming units [CFU]/cm^2^) recovered at regular intervals for 30 days before (open circles) and after (closed circles) the application of SAQAS-A to benches (a/c/e) and glass areas (b/d/f). Experiments were performed over three trials (Trial 1 [a/b], Trial 2 [c/d], and Trial 3 [e/f]). Dotted lines indicate the upper and lower limits of detection. Blue boxes (labelled ’Acceptable’) denote viable counts below the acceptable level of 2.5 CFU/cm^2^; yellow boxes (labelled ’Unacceptable’) denote viable counts above this level.

### SAQAS-A is not effective for 30 days as a biocidal floor treatment

The microbial burden present on floor surfaces was determined before and after SAQAS-A application (Fig.2 A/C/E). The data show there was no reduction in recovered CFU due to the biocide treatment. In fact, the number of samples in which the microbial burden exceeded the acceptable level of 2.5 CFU/cm^2^ was greater after biocide application (46/63 samples [73%]; 15/21 [Trial 1], 15/21 [Trial 2], and 16/21 [Trial 3]) than before (23/63 samples [37%]; 8/21 [Trial 1], 8/21 [Trial 2], and 7/21 [Trial 3]) (Fig.2 A/C/E).

**Figure 2.**
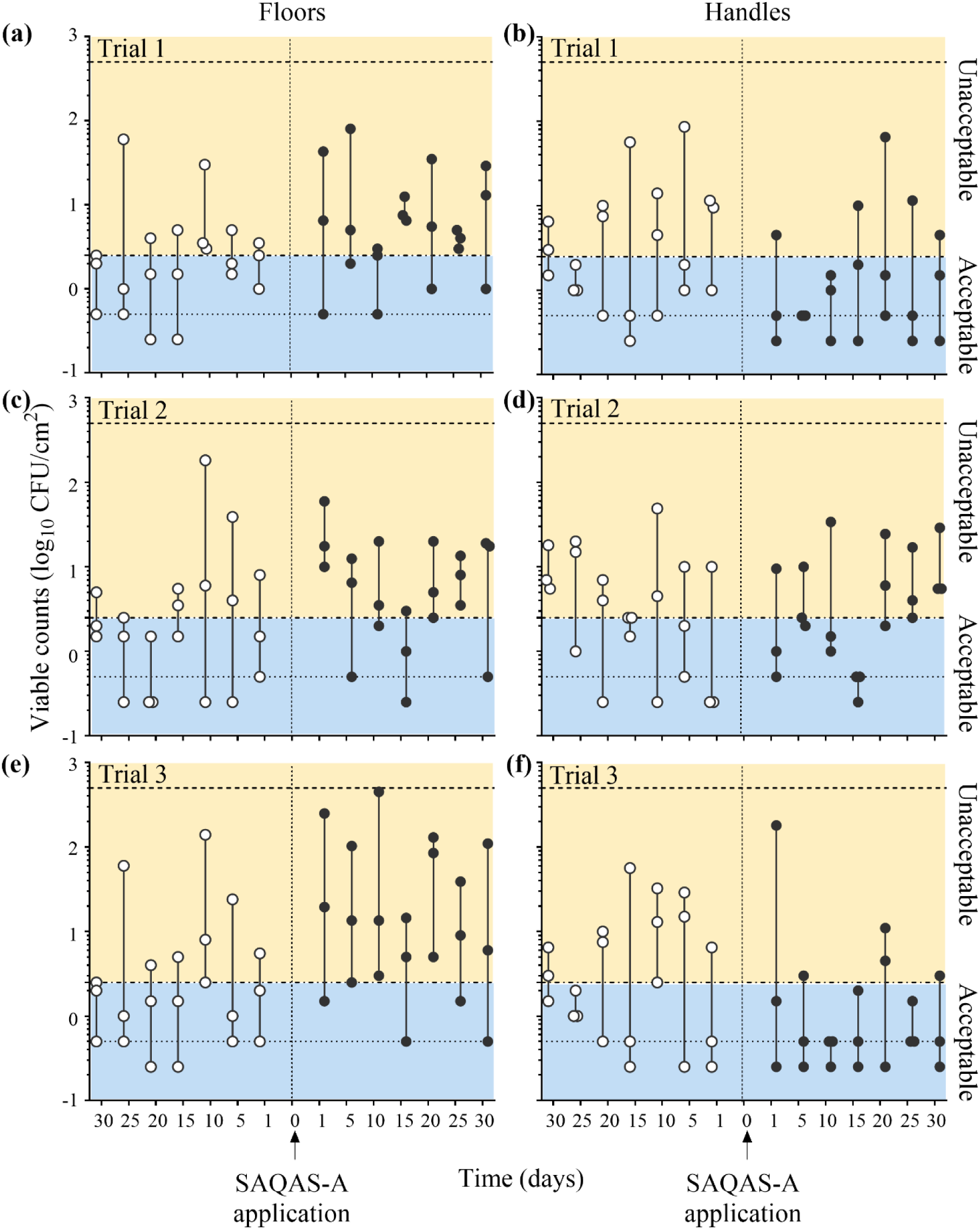
Bacterial counts recovered from floors and handles before and after the application of SAQAS-A. Data presented as individual viable counts (log_10_ colony forming units [CFU]/cm^2^) recovered at regular intervals for 30 days before (open circles) and after (closed circles) the application of SAQAS-A to floors (a/c/e) and handles (b/d/f). Experiments were performed over three trials (Trial 1 [a/b], Trial 2 [c/d], and Trial 3 [e/f]). Dotted lines indicate the upper and lower limits of detection. Blue boxes (labelled ’Acceptable’) denote viable counts below the acceptable level of 2.5 CFU/cm^2^; yellow boxes (labelled ’Unacceptable’) denote viable counts above this level.

### SAQAS-A may offer some advantage as a biocidal treatment of high-touch incubator handles

The microbial burden present on high-touch incubator handles was determined before and after SAQAS-A application (Fig.2 B/D/F). The data show that these high-touch surfaces frequently exceeded the highest acceptable level of 2.5 CFU/cm^2^ during the 30-day period prior to each trial (31/63 samples [49%]; 10/21 [Trial 1], 11/21 [Trial 2], and 10/21 [Trial 3]) (Fig.2 B/D/F). However, during the testing of SAQAS-A, for 2/3 trials, the number of samples exceeding the highest acceptable level halved (10/42 samples [24%]; 5/21 [Trial 1] and 5/21 [Trial 3]) (Fig.2 B/F). For Trial 2, the number of samples exceeding the acceptable level was similar before and after treatment (11/21 and 10/21, respectively) (Fig 2. D). This suggested that SAQAS-A may offer some advantages as a biocidal treatment for high-touch surfaces. However, closer inspection of the data obtained from each individual handle (Fig.3), showed only a marginal benefit that varied by handle and Trial. For example, in Trial 1, the most contaminated handle before the application of SAQAS-A remained so after application (Fig. 3A). In Trial 3, there was a reduction in CFU for the most contaminated handle after SAQAS-A application, though bacterial levels remained above the allowable limit on 3/7 sampling times.

**Figure 3.**
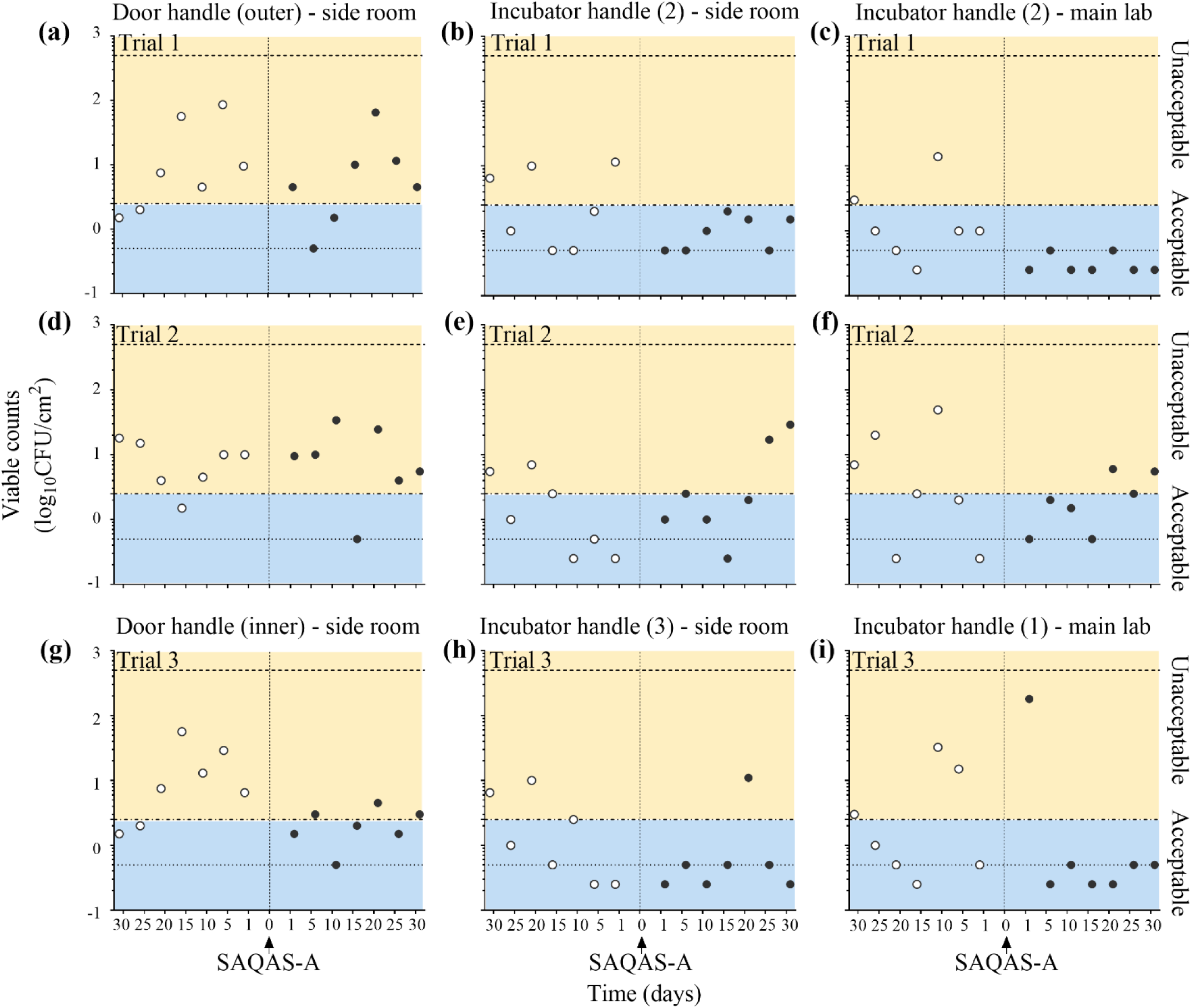
Bacterial counts recovered from handles before and after the application of SAQAS-A. Data presented as individual viable counts (log_10_ colony forming units [CFU]/cm^2^) recovered at regular intervals for 30 days before (open circles) and after (closed circles) the application of SAQAS-A to side room outer door handle (a/d), side room inner door handle (g), side room incubator 2 handle (b/e), side room incubator 3 handle (h), main lab incubator 2 handle (c/f), and main lab incubator 1 handle (i). Experiments were performed over three trials (Trial 1 [a/b/c], Trial 2 [d/e/f], and Trial 3 [g/h/i]). Dotted lines indicate the upper and lower limits of detection. Blue boxes (labelled ’Acceptable’) denote viable counts below the acceptable level of 2.5 CFU/cm^2^; yellow boxes (labelled ’Unacceptable’) denote viable counts above this level.

### Statistical analysis of microbial burden data obtained during real-world testing of SAQAS-A

The practicalities of testing SAQAS-A in a working laboratory meant that trials were performed at different times of the year and using different sampling sites. We analysed the microbial burden data using two Generalised Linear Mixed Models (GLMM) to investigate whether there were any differences in microbial burden before and after the application of SAQAS-A for any of the different types of surfaces and between trials. We also tested for interactions between these variables. For model 1, we compared all data obtained from the pre- SAQAS-A application with all data obtained post-application (Tables 1 and 2). For model 2, we analysed the data obtained over the first 5 days post-SAQAS-A application separately from the data obtained after 5 days post-application (Tables 1 and 2).

**Table 1.** Model statistics.

| Statistical model | F statistic | DF<sub>1</sub> | DF<sub>2</sub> | Significance |
| --- | --- | --- | --- | --- |
| **Model 1** |  |  |  |  |
| Corrected model | 52.25 | 10 | 486 | <0.001 |
| Trial (n=3) | 9.84 | 2 | 486 | <0.001 |
| Surface (n=4) | 20.16 | 3 | 486 | <0.001 |
| Intervention (SAQAS-A application) | 10.25 | 1 | 486 | 0.001 |
| Trial × surface | 4.62 | 6 | 486 | <0.001 |
| Trial × intervention | 3.35 | 2 | 486 | 0.036 |
| Surface × intervention | 79.77 | 3 | 486 | <0.001 |
| **Model 2** |  |  |  |  |
| Corrected model | 2100.11 | 11 | 480 | <0.001 |
| Trial (n=3) | 9.52 | 2 | 480 | <0.001 |
| Surface (n=4) | 58.15 | 3 | 480 | <0.001 |
| Intervention (SAQAS-A application) | 6.17 | 2 | 480 | 0.002 |
| Trial × surface | 6.95 | 6 | 480 | <0.001 |
| Trial × intervention | 3.72 | 4 | 480 | 0.005 |
| Surface × intervention | 301.36 | 6 | 480 | <0.001 |

Key: DF, degrees of freedom

**Table 2.**
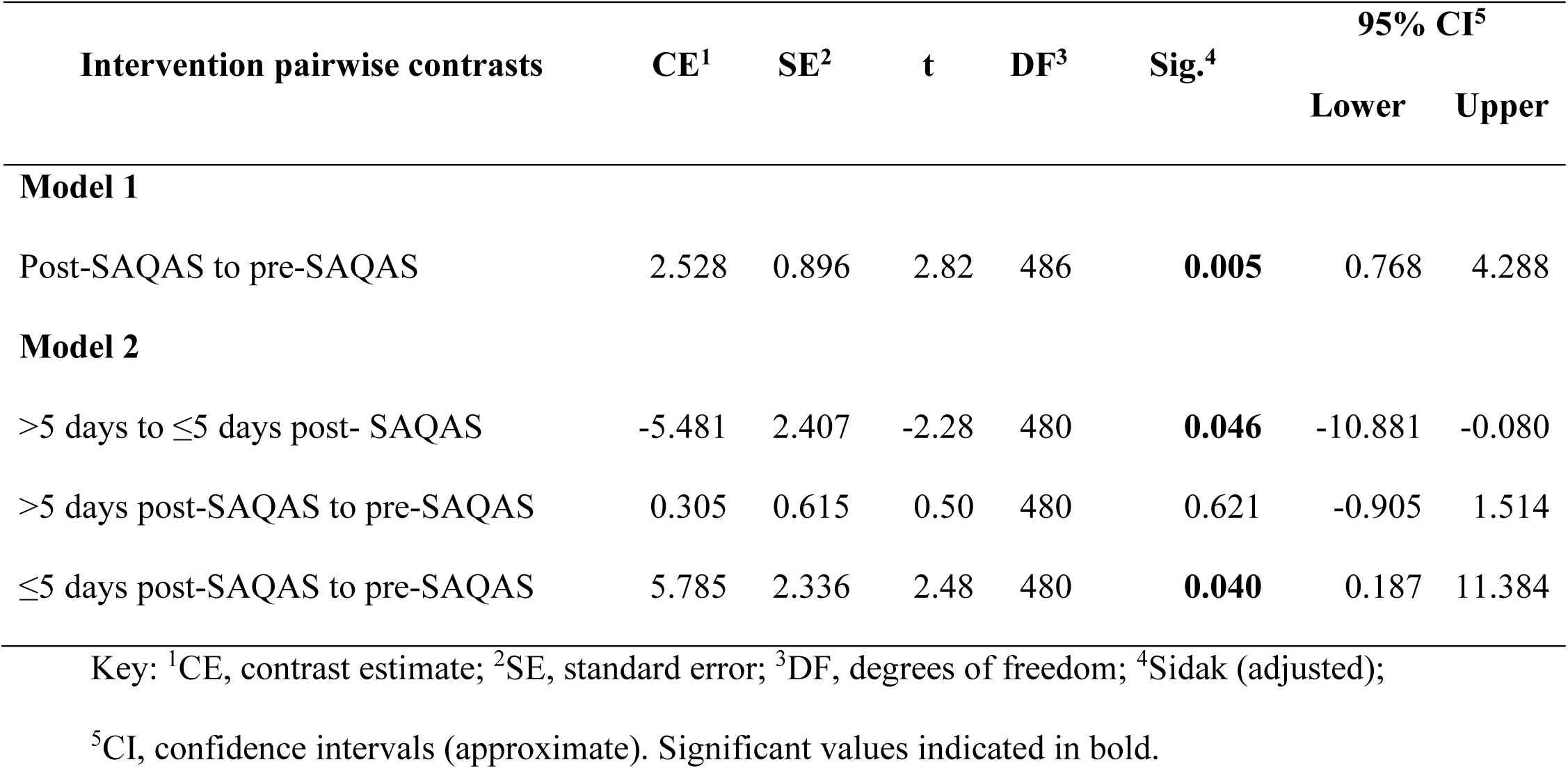
Pairwise contrasts.

The analyses show significant differences between microbial burdens between surfaces, trials, pre-and-post-SAQAS-A application, and two-way interactions between these variables (Tables 1 and 2). However, these differences were indicative of SAQAS-A application having no antimicrobial effect. Model 1 shows that, overall, microbial counts were significantly higher after applying SAQAS-A than before (p=0.005, GLMM with Šidák correction for multiple analyses) (Table 2). Model 2 shows that microbial burdens were significantly higher within the first 5 days after SAQAS-A application (p=0.040, GLMM with Šidák correction for multiple analyses) but then decreased back to pre-intervention levels (p=0.621, GLMM with Šidák correction for multiple analyses) (Table 2).

### Identification of environmental bacteria collected from laboratory surfaces

We selected for identification bacterial isolates recovered from surfaces before and after SAQAS-A treatment based on their dominance on the enumeration plates. Bacterial isolates were stored as frozen stocks and recovered onto Tryptic Soy Agar (TSA) for identification using a mass spectrometry approach. The identified organisms are listed in Table 3. The results show that more Gram-positive species were isolated than Gram-negative (18 and 8, respectively). Of the 8 Gram-negative species isolated, 2/8 were isolated from floors before the application of SAQAS-A and 6/8 were isolated after SAQAS-A application (Table 3). The only Gram-negative bacterium isolated from handles was *Pseudomonas chlororaphis*, after SAQAS-A application. Three Gram-negative bacterial species were isolated from laboratory benches: *Pseudomonas oryzihabitans* before and after SAQAS-A application, *Pseudomonas putida* before SAQAS-A application, and *Roseomonas mucosa* after SAQAS-A application. Only *Ps. putida* has been used by researchers within the laboratory. Of the 18 Gram-positive species isolated, three were *Bacillus* spp. and six were *Staphylococcus* spp. Like the Gram-negative bacteria, Gram-positive bacteria were more likely to be isolated after SAQAS-A application than before (Table 3). More species were isolated from floors and handles than from benches. Of the *Staphylococcu*s spp. 5/6 were isolated from handles post-SAQAS-A application compared to 3/6 from floors and benches (Table 3).

**Table 3.** Environmental microbes recovered during this study.

| Bacterium | Description | Benches |  | Floors |  | Handles |  |
| --- | --- | --- | --- | --- | --- | --- | --- |
|  |  | Pre | Post | Pre | Post | Pre | Post |
| **Gram-negative:** |  |  |  |  |  |  |  |
| *Brucella anthropi* | Formerly *Ochrobactrum anthropi*; opportunistic pathogen(48, 49). | - | - | - | + | - | - |
| *Pseudomonas* spp.: |  |  |  |  |  |  |  |
| *Ps. chlororaphis* | Plant growth-promoting bacterium; also used as a biocontrol agent(50, 51). | - | - | - | - | - | + |
| *Ps. oryzihabitans* | Environmental bacterium; opportunistic pathogen(52, 53). | + | + | - | + | - | - |
| *Ps. putida* | Saprophytic soil bacterium; model research organism(54). | + | - | - | + | - | - |
| *Ps. stutzeri* | Plant growth-promoting bacterium; opportunistic pathogen(55, 56). | - | - | - | + | - | - |
| *Rhizobium radiobacter* | Formerly *Agrobacterium radiobacter*; plant pathogenic bacterium and opportunistic human pathogen(57, 58). | - | - | + | - | - | - |
| *Roseomonas mucosa* | Commonly isolated from the human skin microbiota; used topically to treat patients with atopic dermatitis(59, 60). | - | + | - | + | - | - |
| *Sphingomonas melonis* | Plant pathogenic bacterium(61). | - | - | + | - | - | - |
| **Gram-positive:** |  |  |  |  |  |  |  |
| *Bacillus cereus* group | Complex of closely related lineages with pathogenic potential in humans and animals(62, 63). | + | - | + | + | - | - |
| *Bacillus altitudinis/pumilus* complex | Environmental bacterium; strain with antibacterial properties previously isolated from a laboratory(64, 65). | - | - | + | - | - | - |
| *Brevibacterium casei* | Environmental bacterium reported isolated from cheese and milk; rare opportunistic human pathogen(66, 67). | - | - | - | - | + | - |
| *Brevundimonas diminuta* | Formerly *Ps. diminuta*; environmental bacterium and emerging opportunistic human pathogen(68, 69). | - | - | - | + | - | - |
| *Dietzia cinnamea* | Environmental bacterium; rare opportunistic human pathogen(70, 71). | - | - | - | - | + | - |
| *Kocuria palustris* | Environmental bacterium; emerging opportunistic human pathogen(72, 73). | - | - | - | + | - | - |
| *Kocuria rhizophila* | Environmental bacterium; opportunistic human pathogen(72, 74). | - | + | - | + | - | + |
| *Kytococcus sedentarius* | Formerly *Micrococcus sedentarius*; frequently isolated from indoor air; emerging opportunistic human pathogen(75, 76). | - | - | - | + | - | + |
| *Microbacterium paraoxydans* | Pathogen of fish; rare opportunistic human pathogen(77, 78). | - | - | - | - | - | + |
| *Micrococcus luteus* | Environmental bacterium also found as part of human skin microbiota; opportunistic human pathogen(79, 80). | - | - | - | + | - | - |
| *Paenibacillus pabuli* | Formerly *Bacillus pabuli*; environmental bacterium which has been found contaminating multi-use tear drops, gels, and ointments(81, 82). | - | + | + | - | + | + |
| *Peribacillus simplex* | Formerly *Bacillus simplex*; environmental bacterium with plant-growth-promoting, biocontrol, and bioremediation properties(83, 84). | - | - | - | + | - | - |
| *Staphylococcus* spp.:<br>*Staph. capitis* | Member of human skin microbiota; opportunistic human pathogen(85, 86). | - | + | - | + | + | + |
| *Staph. epidermidis* | Member of human skin and mucous membrane microbiota; opportunistic human pathogen (most common cause of implant-associated infections)(87). | - | - | + | - | - | + |
| *Staph. hominis* | Member of human skin microbiota; opportunistic pathogen(88, 89). | - | - | - | + | - | + |
| *Staph. pasteurii* | Environmental bacterium also found in food products; rare opportunistic human pathogen(90, 91). | - | + | - | - | - | - |
| *Staph. saprophyticus* | Environmental bacterium also found as part of genitoanal microbiota; common cause of urinary tract infections(92, 93). | - | - | - | - | + | + |
| *Staph. warneri* | Member of human skin and gut microbiota; opportunistic pathogen(86, 94, 95). | - | + | - | + | - | + |

Key: +, present; -, absent

## Discussion

We have tested a commercially available contact-dependent SAQAS-type biocide formulation that is based on the active siloxane-anchored quaternary ammonium salt (“SAQAS-A”) and is available in an easily applicable spray form. We demonstrate that SAQAS-A was unable to protect surfaces for 30 days, as claimed by the manufacturer(20), in a real-world scenario that included routine chemical disinfection of surfaces.

According to the manufacturer, SAQAS-A becomes active after drying onto a surface, with a single-molecule layer of biocide forming on the surface(20). No special treatment is required for biocide anchoring or attachment, and the product is suitable for use on any surface. They further claim that a single application of biocide provides up to 30 days of stability that can withstand surface cleaning. Our previously published work demonstrated that SAQAS-A treatment protected surfaces in assays using treated glass and low-density polyethylene carriers(35). The lack of activity of the SAQAS-A treatment within a working microbiology laboratory was disappointing, indicating that Gram-positive bacteria, including Staphylococcus spp., survived on SAQAS-A-treated surfaces. This is despite in vitro tests showing effective killing of S. aureus(35). Studies of hospital fomites have found that the predominant microbes are Staphylococcus spp.(36–38). We observed that treated areas attracted more dust than untreated areas, which may explain the higher microbial burdens, as dust particles provide a surface for microbes to adhere to(39–41). We speculate that the survival of environmental bacteria on treated surfaces may be due to their carriage on dust or other particles, which form a barrier between the bacteria and the SAQAS layer. We believe this should be investigated further. We did not follow the transmission of bacteria to and from the surfaces. However, future work may track bacteria found on the hands of workers in contact with treated or untreated surfaces and monitor the transmission of specific isolates, as well as whether surface treatments can reduce the prevalence or transmission rate of specific organisms.

The routine disinfection of treated surfaces may be a factor in the poor long-term protection seen, and we suggest that regular cleaning with liquid disinfectants may erode the protective layer. The international standards by which the potency of antimicrobial surfaces is often measured (for example, the Japanese Industrial Standard (JIS) Z 2801:2012 and International Organisation of Standardisation (ISO) 22196:2011(42)) have not covered the aspect of long-term durability, and so regulatory and consumer protection agencies are restricted in their ability to provide clear guidance for companies wishing to make claims of durability and longevity for biocidal surface applications. New standards that incorporate tests of the durability of antimicrobial treatments, such as PAS 2424:2014(43), are a step in this direction. Indeed, in response to the COVID-19 pandemic, the United States Environmental Protection Agency (EPA) updated its guidance for companies seeking to claim residual activity for their products, as reviewed by Donskey(22). Importantly, this guidance includes a method to simulate cycles of in-service disinfection and cleaning(44). On its website, the EPA lists residual antimicrobial products approved for use against SARS-CoV-2(45). At the time of writing, the list was last updated on April 23, 2024, and contains only two products, both of which are copper-containing paints. Future work should examine whether biocidal layers withstand cleaning and analyse the long-term chemical stability and chemical durability of biocides on surfaces. An assessment of the durability of surface coatings might be a good approach to explain the failure of SAQAS-A in real-world conditions.

## Materials and Methods

Cotton swabs were purchased from Global Science & Technology, Auckland, New Zealand, and sterilised in-house by autoclaving. Microorganisms were released from swabs in either Peptone Water (Fort Richard, Auckland, New Zealand) or Letheen Broth (Merck Millipore Corporation, USA). Microbes were cultured on Difco Tryptic Soy Agar (TSA) and differential media for Enterobacteriaceae (Eosin Methylene Blue Agar [EMBA]) (Fort Richard, Auckland, New Zealand), *Staphylococcus* spp. (Mannitol Salt Agar [MSA]) (Fort Richard, Auckland, New Zealand), *Bacillus* spp. (Mannitol Egg Yolk Polymyxin Agar [MYPA]) (Fort Richard, Auckland, New Zealand), *Pseudomonas* spp. (*Pseudomonas* Selective Agar [CFC]) (Fort Richard, Auckland, New Zealand), and fungi (Potato Dextrose Agar [PDA]) (Fort Richard, Auckland, New Zealand). Strains were stored at -80 °C and recovered on TSA with incubation at 28 °C for 2 days.

A commercially available, spray-on biocide formulation (Zoono Z-71 Microbe Shield Surface Sanitiser) (Elitepac NZ Ltd.) was used and is referred to as SAQAS-A. Surface disinfectants were Trigene Advance (Invitro Technologies, New Zealand), used at a 1:100 dilution in water, and ethanol, used at 70 % by volume in water.

### Environmental sampling

Three high-traffic floor areas, three routinely used bench areas, three frequently touched handles, and three glass surfaces were designated as sampling sites within the PC2 laboratory (Fig. 4). The chosen areas (Table 4) were routinely cleaned according to local laboratory procedures during the trial period. Floors were cleaned twice a week using Trigene Advance, and bench surfaces were cleaned after every period of microbiological work using Trigene Advance, followed by 70 % ethanol. Door handles were cleaned once a week using 70 % ethanol. Incubator handles were cleaned less regularly. There was no routine cleaning of window-glass surfaces. Trigene Advance was applied for a contact time of 10 min and 70 % ethanol until it had evaporated (approx. 1 min). Each designated sampling area was 20 cm^2^. Each area was sampled between 11:30 a.m. and 1:30 p.m. every 5 days over 30-day periods before and after SAQAS-A treatment, and bacterial enumeration on agar plates was used to determine the microbial burden. Experiments were repeated on three separate occasions (Trials 1, 2, and 3).

**Figure 4.**
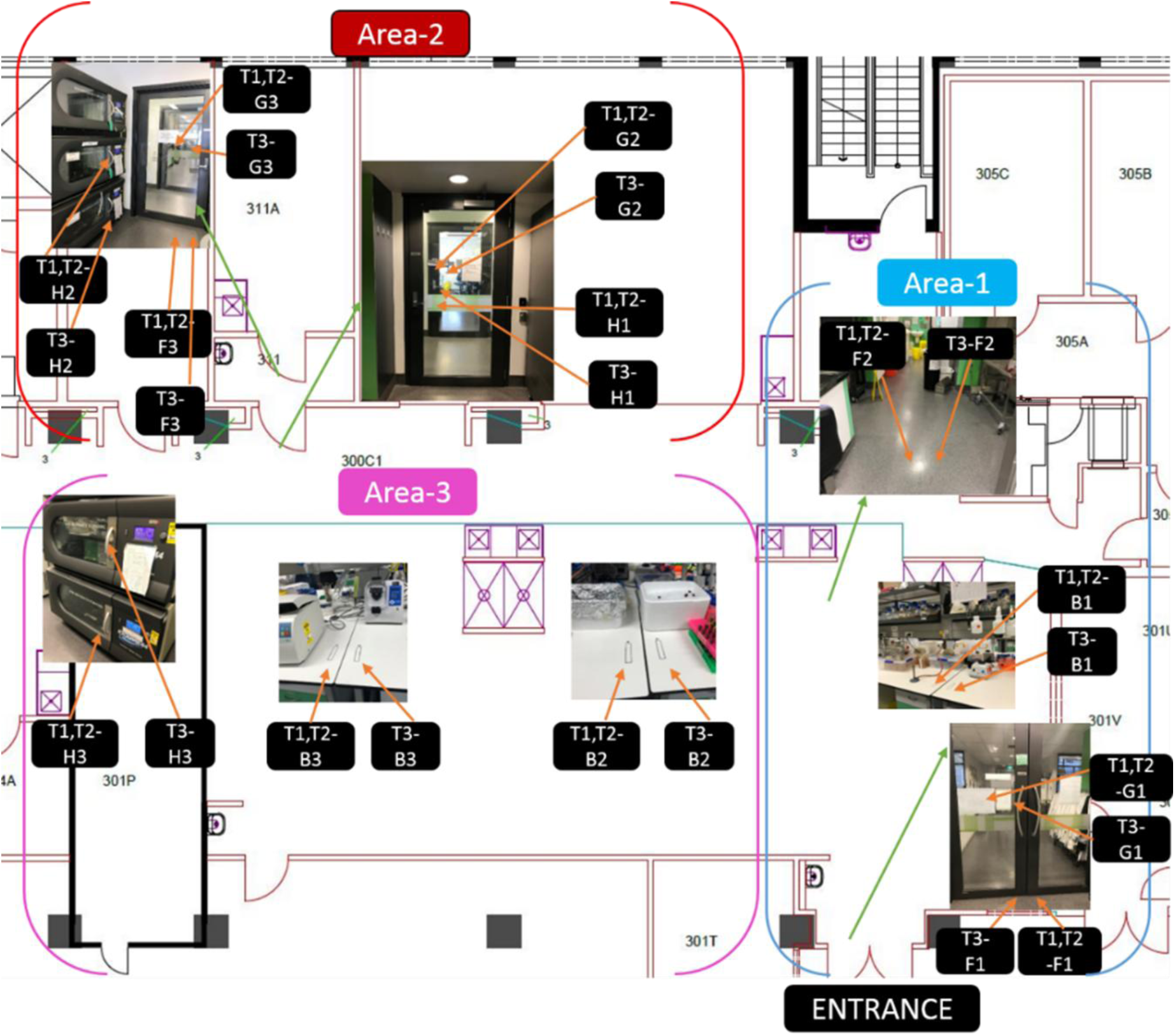
Laboratory areas chosen for real-world testing. The laboratory was divided into three areas as indicated on the plan. Key: B, bench; F, floor; G, glass; H, handle; T1, trial 1; T2, trial 2; T3, trial 3.

**Table 4.** Laboratory areas used for study.

| Surface | Trials 1 and 2 | Trial 3 |
| --- | --- | --- |
| Benches |  |  |
| Area 1 – B1 | Main lab, left | Main lab, right |
| Area 3 – B2 | Main lab, left | Main lab, right |
| Area 3 – B3 | Main lab, left | Main lab, right |
| Floors |  |  |
| Area 1 – F1 | Main lab (entrance), right | Main lab (entrance), left |
| Area 1 – F2 | Main lab, left | Main lab, right |
| Area 2 – F3 | Side room, right | Side room, left |
| Glass |  |  |
| Area 1 – G1 | Main lab (entrance), right | Main lab (entrance), left |
| Area 2 – G2 | Side room (door 1), right | Side room (door 1), left |
| Area 2 – G3 | Side room (door 2), right | Side room (door 2), left |
| Handles |  |  |
| Area 2 – H1 | Side room (door 1), outside | Side room (door 1), inside |
| Area 2 – H2 | Side room, shaking incubator (2) | Side room, shaking incubator (3) |
| Area 2 – H3 | Main lab, shaking incubator (2) | Main lab, shaking incubator (1) |

For sampling, designated areas were swabbed with a sterile cotton swab that had first been moistened in sterile saline (0.85 % w/v NaCl) and then pressed against the inside wall of the tube to remove any excess liquid. Surfaces were swabbed horizontally, vertically, and then diagonally for approximately ten passes in each direction, with the swab being rotated axially between the thumb and forefinger to ensure all sides of the swab were used. Sufficient pressure was applied to the surface through the swab to cause the swab shaft to begin to bend. The swab heads were then placed into a bottle containing 10 mL of peptone water (Trial 1) or 2 mL Letheen broth (Trials 2 and 3), and the shaft was snapped off aseptically against the inner wall of the bottle, leaving the cotton head in the bottle. Letheen broth was used as a precaution to neutralise any biocide also removed from the surface that we speculated would reduce the microbial burden recovered on the pour plates. The capped bottles were vortexed for 3 × 20 s with 20 s of rest between each vortex. Aliquots of 100 µL were plated onto TSA, incubated at 28 °C for 2 days, and CFU counted. In Trial 1, an aliquot of 200 µL was plated onto differential media (EMBA, MSA, MYPA, CFC, and PDA) and incubated at 28 °C for 2 to 4 days. A selection of the predominant colony types was purified from sampling plates and identified by MALDI-TOF at LabPlus, Auckland.

### Generalised Linear Mixed Models (GLMM)

The data obtained from testing the real-world efficacy of SAQAS-A on laboratory surfaces were analysed using two Generalised Linear Mixed Models (GLMM) using IBM SPSS Statistics for Windows (Version 26.0) (IBM Corp., Armonk, NY). A mixed model was used because multiple measurements were taken from the same areas, and this type of model considers the measurements within each area as being related(46). The models incorporated a negative binomial distribution with logarithmic links and repeated measures. A Sidak sequential adjustment was applied to all post hoc pairwise comparisons to adjust for multiple testing(47). Factors included in model 1 were intervention (pre/post-SAQAS-A treatment), surface type (bench/floor/glass/handles), and trial (1–3). The same factors were included in model 2, but this model also tested the strength of the intervention in the first 5 days (post ≤5 days) versus longer-term (post >5 days: 10-to-30 days).

## Acknowledgements

The New Zealand Ministry of Business, Innovation and Employment (MBIE) funded this project through the Biocide Tool Box (BTB) research program (grant number UOAX1410). The funder had no role in study design, data collection and interpretation, or the decision to submit the work for publication.

## Transparency declarations

SS has provided expert opinion to the New Zealand Commerce Commission regarding Zoono Ltd products. All other authors: none to declare.

## Data availability

The data underlying this article are available in the article (Table 3) and in Figshare (accessed using https://figshare.com/s/fd3c3511fcc2aadb3e30 [currently set to private but will be published on acceptance]).

